# Reconstructing phylogeny of the hornless rhinoceros Aceratheriinae

**DOI:** 10.1101/2022.02.04.478979

**Authors:** Xiaokang Lu, Tao Deng, Luca Pandolfi

**Affiliations:** Human Anatomy and Histology Department, Henan University of Chinese Medicine, Zhengzhou 450008, China; Key Laboratory of Vertebrate Evolution and Human Origins, Institute of Vertebrate Paleontology and Paleoanthropology, Chinese Academy of Sciences (CAS), Beijing 100044, China; State Key Laboratory of Palaeobiology and Stratigraphy, Nanjing Institute of Geology and Palaeontology, Chinese Academy of Sciences (CAS), Nanjing 210008, China; CAS Center for Excellence in Life and Paleoenvironment, Beijing 100044, China; University of Chinese Academy of Sciences, Beijing 100049, China; A Dipartimento di Scienze della Terra, Università di Firenze, Firenze 50121, Italy

## Abstract

This study presents the first phylogenetic analysis focused on the Subfamily Aceratheriinae to date, with 391 characters coded from 43 taxa at the species level. We added 77 newly defined and 33 revised characteristics, including features of the skull, teeth, and postcranial bones. In the present analysis, the tribe Teleoceratini, as well as the tribe Aceratheriini, was reclassified within Aceratheriinae, however, the traditionally established monophyly of each tribe was decomposed. Combined with detailed morphological comparisons, we reconstructed the phylogeny of Aceratheriinae and revised the diagnosis of Aceratheriinae. The reported skull and teeth specimens of *Turkanatherium* from the late Early Miocene indicate that it is not an acerathere and has been placed as a basal rhinocerotid. The Aceratheriinae has undergone evolutionary adaptation several times during the early stage of evolution, and several genera from the Oligocene to the Early Miocene have been reconstructed as early diverging taxa, such as *Molassitherium, Protaceratherium, Plesiaceratherium* and *Chilotheridium*; *Aprotodon* and *Mesaceratherium* from the Late Oligocene to the Early Miocene were united as the earliest divergent clade of Aceratheriinae; meanwhile, due to the limited materials, the phylogenies of *Floridaceras* and *Dromoceratherium* are unstable in this analysis, and both have been tentatively considered as the early diverging clades of Aceratheriinae. *Alicornops* was reclassified a member of Teleoceratini. Aceratheriini and Teleoceratini have been redefined as two highly specialized groups. Furthermore, Aceratheriini has been divided into two/three clades based on the difference in the skull outlines and the occlusal patterns of the cheek teeth.

## INTRODUCTION

Aceratheres are an extinct group of rhinocerotids widespread throughout Eurasia, North America and Africa during the Neogene, characterized by the absence of the nasal and/or frontal horns. The systematic position and the phylogenetic relationships of several genera and species identified as aceratheres remain debated or poorly known.

The first hornless genus *Aceratherium* from the Late Miocene was recorded by Kaup (1832) in Eppelsheim, Germany. Based on this record, together Dollo (1885) established the subfamily Aceratheriinae as a member of the Family Rhinocerotidae, which has another two subfamilies Rhinocerotinae and Elasmotheriinae (Prothero et al., 1989; Heissig, 1999; Antoine, 2002). Osborn (1900) recognised the subfamily Aceratheriinae and included within it some Oligocene and Miocene European species, giving the following diagnosis: “dolichocephalic with long, narrow nasals; smooth or with rudimentary horns at sides of the tips; frontals finally developing horns; large cutting teeth; relatively persistent tetradactyl manus; long-limbed” (Osborn, 1900: 240). Osborn (1900) included the following species within Aceratheriinae: *Aceratherium filholi, A. lemanense, A. platyodon, A. blanfordi, A. tetradactylum, A. incisivum*. Heissig (1973) included within Aceratheriinae the tribes Teleoceratini Hay, 1902 which includes the brachypotheres *Teleoceras* Hatcher, 1894, *Brachypotherium* Roger, 1904, *Aprotodon* Forster-Cooper, 1915 and *Diaceratherium* Dietrich, 1931, and Aceratherini Dollo, 1885; the latter includes the genera *Aceratherium* Kaup, 1832, *Plesiaceratherium* Young, 1935, *Chilotherium* Ringström, 1924, *Aphelops* Cope, 1873. In 1989, Heissig includes within Aceratherinii the genera *Mesaceratherium* Heissig, 1969, *Alicornops* Ginsburg & Guérin, 1979, *Aceratherium* Kaup, 1832, *Plesiaceratherium* Young, 1935, *Hoploaceratherium* Ginsburg & Heissig, 1989, *Aphelops* Cope, 1873, *Peraceras* Cope, 1880, *Chilotheridium* Hooijer, 1971, *Turkanatherium* Deraniyagala, 1951, *Chilotherium* Ringström, 1924, *Subchilotherium* Heissig, 1972, *Acerorhinus* Kretzoi, 1942.

In the following two hundred years, there have been 90 species referred to Aceratheriinae but According to Prothero (2005), this taxon has been used as a taxonomic wastebasket for all hornless rhinoceroses despite some taxa identified as aceratheres may have had small horns. Not all hornless rhinoceros could be classified as Aceratheriinae, which is advanced and includes rhinoceros with a small nasal horn. Other features that characterize Aceratheriinae include a deep nasal notch, reduced upper incisor I1, considerably large lower incisor i2, brachycephalic skull, tetradactyl manus, and shortened and massive metapodials (Heissig, 1989; Cerdeño, 1995).

The definition of Aceratheriinae has been in dispute for many years. Pavlow (1892) united *Aphelops, Teleoceras* and *Brachypotherium*, but assigned *Aceratherium* to the subfamily Rhinocerotinae. Scott and Osborn (1898) recognized the close relationship between *Peraceras* and *Aphelops*. Heissig (1973, 1999) is the first to include the tribe Teleoceratini in Aceratheriinae to this subfamily. Prothero et al. (1986, 1989) suggested that only the members of Aceratheriini could be recognized as Aceratheriinae; thus, the tribe Teleoceratini should be member of Rhinocerotinae. Using 72 characters coded from 43 rhinocerotid taxa, Cerdeño (1995) noted that Teleoceratini is more closely related with Aceratheriini than the other groups within Rhinocerotidae, but this study has yet resolved the phylogeny of several genera previously referred to Aceratheriinae, such as *Protaceratherium, Mesaceratherium*, and *Aprotodon* (Fig. 1).

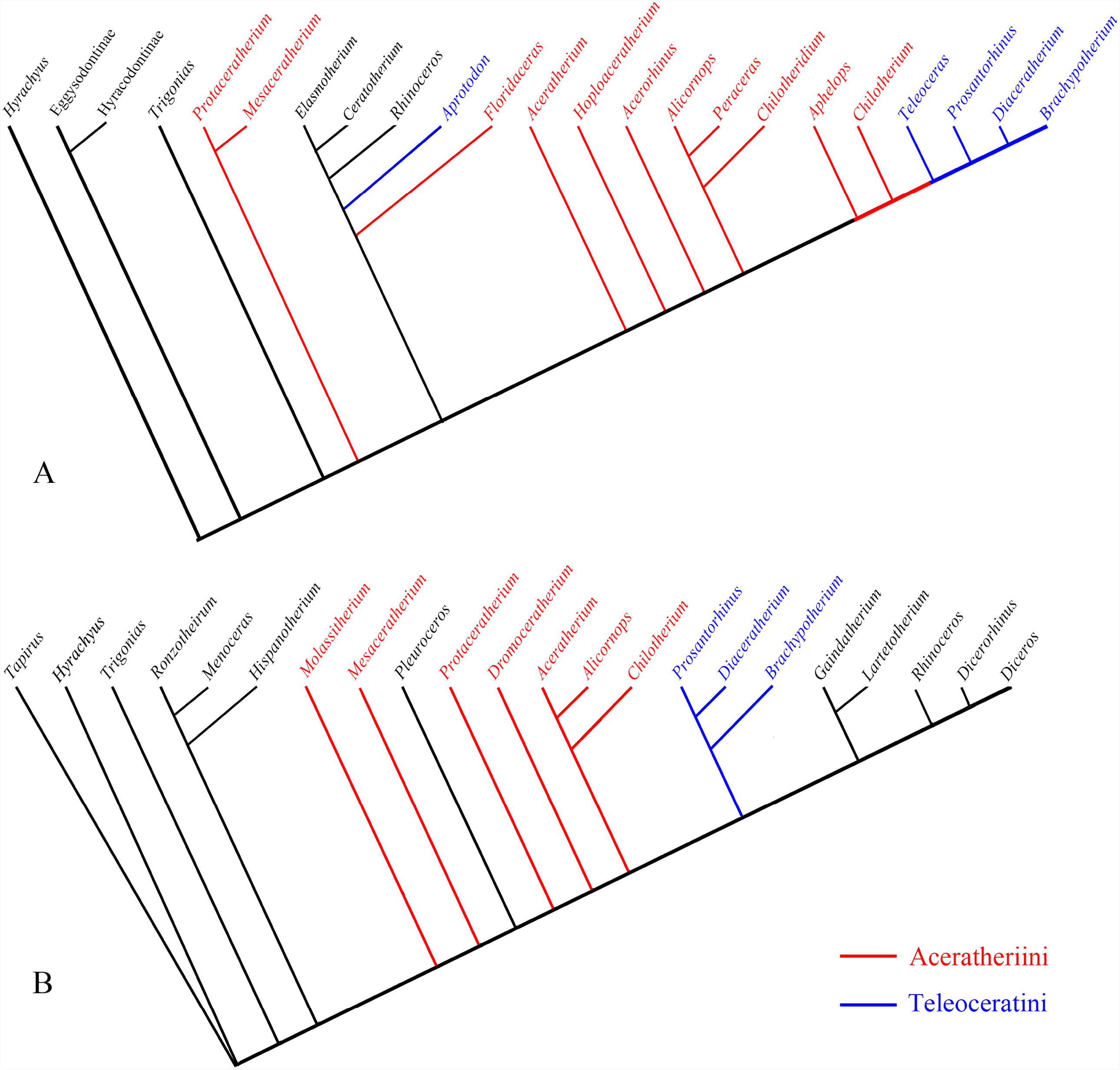

In order to explore the phylogeny of the elasmotheres, Antoine (2002) expanded the matrix to 282 characters, including all the details of the morphology of this group.

Deng (2008) successfully used this matrix to discuss the taxonomic identity of new elasmotheres. However, this combination of characters could not ‘resolve the phylogeny of the acerathere (Antoine et al., 2003, 2010) (Fig. 1), despite the addition of several features by Lu (2013).

Aceratheriinae (Aceratheriini or Aceratheriina) represents a monophyletic group within the subfamily Rhinocerotinae according to Antoine et al. (2002), Antoine (2003), Antoine et al. (2010), Becker et al. (2013) and Pandolfi (2015). Aceratheres are included within the subfamily Aceratheriinae by Prothero (2005) and also by Heissig (2012), who consider this group as sister to Rhinocerotinae + Elasmotheriinae clade.

In the latest analysis of rhinocerotids, three subfamilies, including Aceratheriinae, Rhinocerotinae, and Elasmotheriinae have been reclassified as polyphyletic groups, with confusing relationships (Lu et al., 2016).

This study aims to produce a new inclusive cladistic analysis of Aceratheriinae, for both taxa and characters, partly including previously defined characters. We also aim to reappraise the definition of Aceratheriinae and clarify the taxonomic position of several genera based on both the traditional taxonomic opinion and the results of the present cladistic analysis. The analysis and discussion are based on the previously established phylogenetic relationship of Aceratheriinae and the character polarities of primitive or advanced.

## MATERIALS AND METHODS

After an extensive examination of the morphology of the members of Rhinocerotidae, we added 77 newly defined characters, and revised 33 previously used characters, covering features of the skull, teeth, and postcranial bones. In total, 391 characters were used in this analysis, including 92 characters of cranial, 168 of teeth, and 131 of postcranial (SIs. 1, 4).

Because there is no consensus on the phylogeny of Aceratheriinae, the present analysis incorporated all the genera once referred to as Aceratheriini and Teleoceratini, including 24 taxa, and other genera of rhinocerotoids, including 18 taxa, which provided a larger basis for determining the character polarity of Rhinocerotidae. The outgroup includes four taxa, *Hyrachyus modestus*, which is the most completely identified species within the genus. All the taxa in this analysis are at a species level, represented by the type species of each genus. *Teleoceras* is known for graviportal limb, but the concerning materials are at present not accessible for the type species *Teleoceras major*, accordingly we further included the most characterized species *Teleoceras fossiger*. The available specimens of the type species of *Aprotodon* include a crushed skull and teeth fragments; thus, we chose the Chinese species *Aprotodon lanzhouensis*, the phylogenetically parts of which body are well known, particularly the lower incisor i2. For the same reasons, we included two taxa of *Hispanotherium*, and two extant taxa of *Rhinoceros*. The character scoring relied on direct observation or bibliography data (SI. 2).

The analysis was performed using TNT version 1.5 (Goloboff et al., 2008; Goloboff and Catalano, 2016), with 1000 replicates, tree bisection-reconnection (TBR) branch swapping, and 100 trees saved per replicate. All characters were equally weighted and unordered. The Bremer support values were calculated in TNT by running a traditional search on the most parsimonious tree, with 1000 replications and TBR branch swapping.

## RESULTS

The maximum parsimony analysis performed by TNT resulted in six trees in 1,620 steps (CI = 0.31, RI = 0.51). A lower CI value indicates a high level of homoplasy. A strict consensus tree and a 50% major consensus tree were constructed, and the latter recovered an overall well-resolved topology of Aceratheriinae (Fig. 2). The Bremer support values are low for many clades. This is likely due to the high proportion of missing data and homoplastic characters, which could increase the taxonomic instability in the suboptimal tree, thereby causing the clades to collapse. In the following discussion, the synapomorphy list of each node and the determination of the clade are based on the 50% major consensus tree (SI. 5). The reconstructed tree does not follow the widely recognized opinion regarding the phylogeny of Rhinocerotinae and Elasmotheriinae (Cerdeño, 1995; Antoine, 2002; Antoine et al., 2010). The taxa of Elasmotheriinae and Rhinocerotinae were reclassified as a major polyphyletic clade, and, their relationship will not be discussed here.

In both the strict consensus tree and the 50% majority tree, all taxa that were previously referred to the tribe Aceratheriini and the tribe Teleoceratini were clustered to a large clade (Fig. 2, Node A), except for *Turkanatherium*. Seven characters were presented to support this node: the palatine fissure extends backward toward the posterior part of the diastema (Cha. 2^0-1^); the infraorbital foramen is located below the nasal notch (Cha. 24^0-1^); the ventral rim of the mandible is convex (Cha. 88^0-1^); both the second lower incisors (i2) are greatly divergent (Cha. 106^0-1^); the labial wall of the lower cheek teeth shows a vertical wrinkle in the enamel (Cha. 200^0-1^); and the labial cingula of the lower premolar are reduced to residuals (209^0-1^).

In the present analysis, *Aprotodon* and *Mesaceratherium* were united in the earliest divergent clade of Aceratheriinae, consistent with their mosaic characters, such as less advanced upper premolars, greatly specialized lower incisor (i2) and symphysis (Deng, 2013).

The results of the present analysis did not support the previously established monophyly of Aceratheriini (Prothero and Schoch, 1989; Heissig, 1989; Cerdeño, 1995). *Turkanatherium* has been precluded from Aceratheriinae. The rest of the 17 taxa, except *Mesaceratherium*, which is in a clade with *Aprotodon* in both the strict and 50% majority consensus trees, have been decomposed into seven single genus clades and three small clades (Fig. 2). *Alicornops* has long been referred to Aceratheriini (Prothero et al., 1986; Cerdeño, 1995), but is placed as a sister group to the clade merging four genera of Teleoceratini in the 50% majority tree of the present analysis (Fig. 2) (Node C, Cha. 95^1-0^; 109^1-0^, 154^2-1^, 212^1-2^, 314^1-0^, 360^1-0^).

*Hoploaceratherium, Aphelops*, and *Acerorhinus* form a small clade with a low Bremer support value (Node D, Char. 33^1-0^, 42^0-1^, 120^0-1^, 125^2-1^, 182^0-1^, 211^1-3^, 292^0-1^, 359^1-0^). Another small clade clustering four Late Miocene genera, including *Aceratherium, Peraceras, Chilotherium*, and *Shansirhinus*, represented the most advanced group of Aceratheriini, with high Bremer support value (Node E, Char. 26^0-1^, 42^0-1^, 56^0-1^, 60^0-1^, 183^0-1^, 346^1-0^). One unexpected clade is *Galushaceras* and *Subchilotherium*, both of which are from the Middle Miocene (Heissig, 1972; Prothero, 2005) (Node F, 132^1-0^, 154^2-3^, 164^1-0^, 184^0-1^).

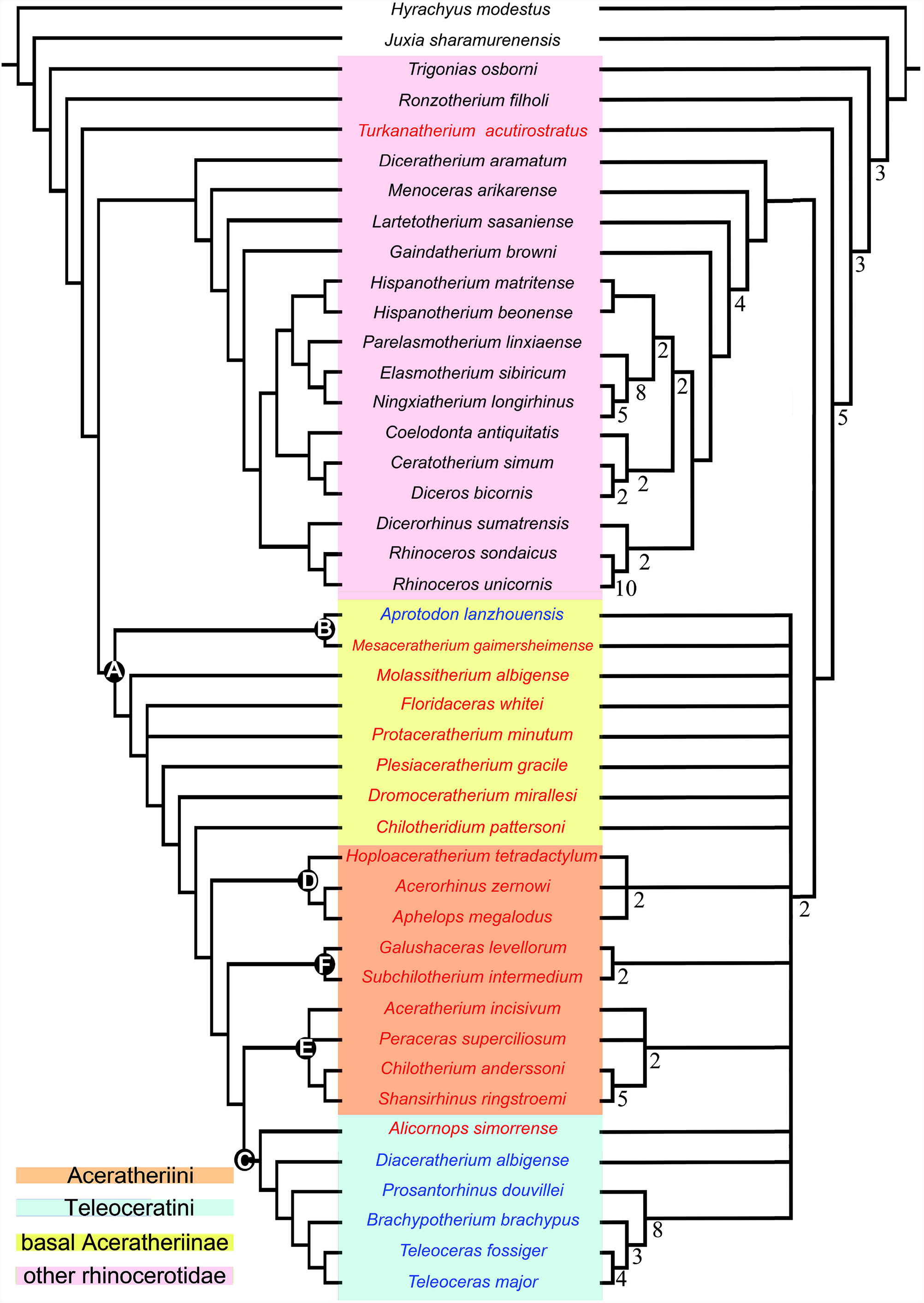

## DISCUSSION

### Diagnosis of Aceratheriinae

The phylogenetic analysis here provided a set of new characters those would be used as diagnostic features. On the other hand, however, some features that have long been considered as diagnostic of Aceratheriini and Teleoceratini were not present as synapomorphies of node A, including the retracted nasal notch, tusk i2, fully reduced I1, deeply constricted lingual cusps, brachycephalic skull, and graviportal limbs. These features demonstrate a gradually changing tendency, and the maximum only occurs in limited taxa, such as *Chilotherium* and *Teleoceras*.

The clear starting point when studying the characteristics of hornless rhinoceros is the horn. A small horn and the enlarged upper incisors I1 are two crucial characteristics that Prothero et al. (1986) used to include Teleoceratini to Rhinocerotinae rather than Aceratheriinae. However, among members once classified as Aceratheriini, at least four genera developed a small nasal horn, including *Chilotheridium, Hoploaceratherium, Peraceras*, and *Shansirhinus*. In Teleoceratini, the presence of a nasal horn has an interspecies variation in *Diaceratherium* and *Brachypotherium* (Répelin, 1917; Heissig, 1999, 2017) (Fig. 3). Nasal horns had developed in different lineages of rhinocerotids several times, as well as the great enlargement of the upper incisors I1, which has occurred in *Mesaceratherium* since the Oligocene (Heissig, 1969).

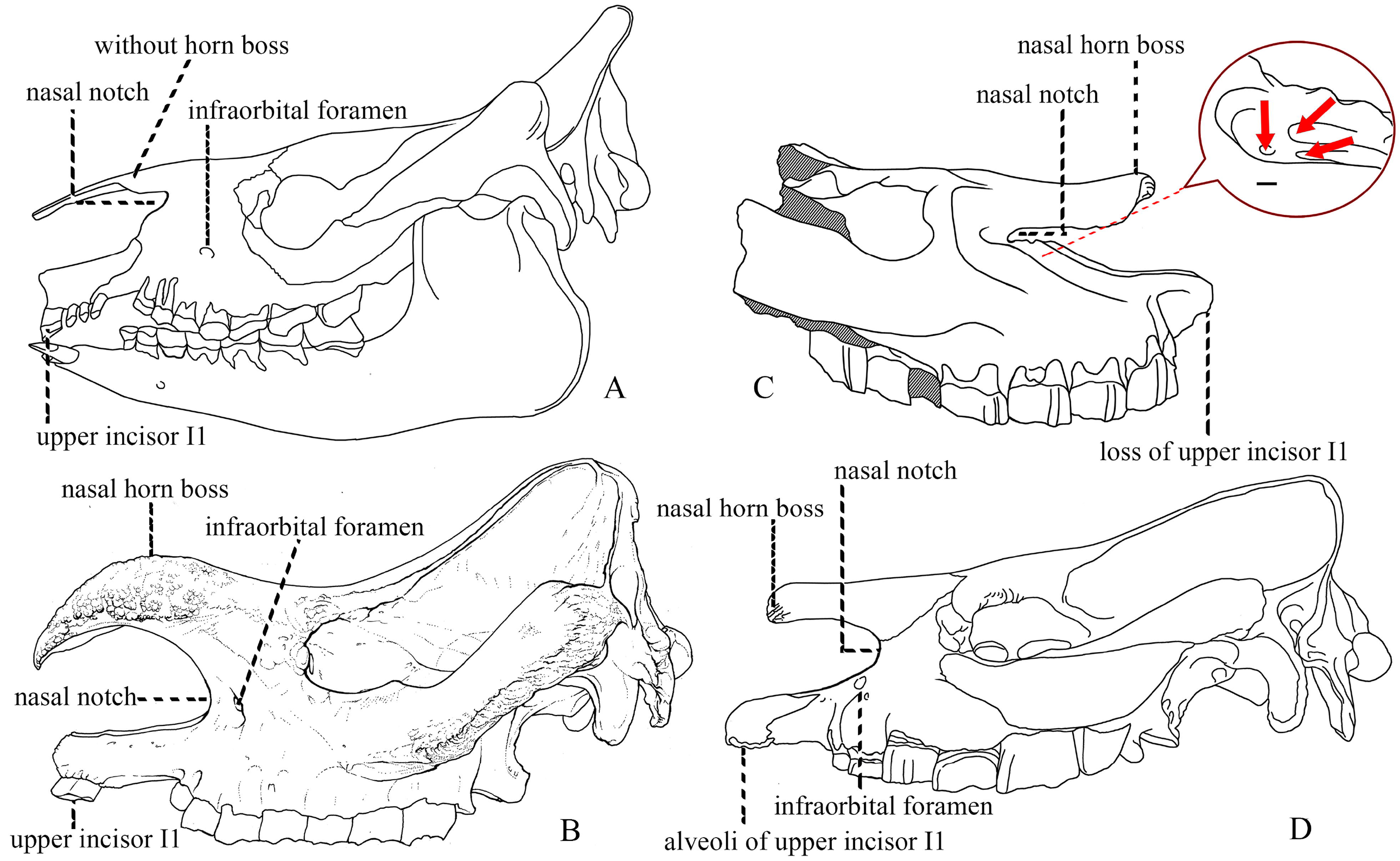

Another notable structure lies in the rostral end of the skull and involves two aspects, namely the retracted nasal notch and the specialized incisors, both of which are closely related to feeding behavior (Prothero et al., 1986; Heissig, 1989, 1999; Cerdeño, 1995). The enlarged lower incisor (i2) is a typical feature differentiating rhinocerotids from other rhinocerotoids and were maximized in the evolution of Aceratheriini and Teleoceratini, exemplified by *Chilotherium* and *Teleoceras*, respectively (Ringström, 1924; Deng, 2001; Prothero, 2005) (Fig. 4). However, the development of the upper incisor I1 diverged in the two tribes; Teleoceratini retained the upper incisor I1, while in the advanced members of Aceratheriini, I1 has been lost, and the enlarged lower incisor i2 subsequently loses the occlusal surface on the internal edge (Fig. 3, 4) and developed a labially upturning occlusal surface, such as in *Chilotherium* (Deng, 2001; Lu, 2013) (Fig. 4).

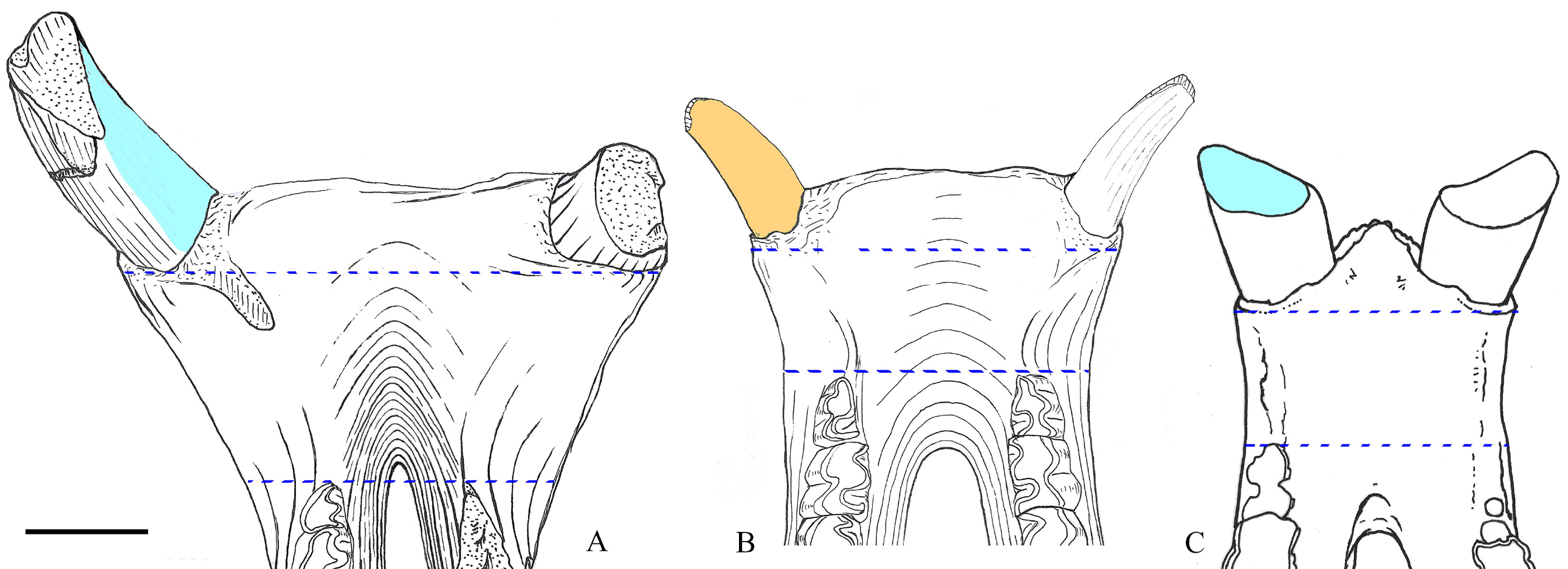

The retracted nasal notch is a gradually evolving structure that has occurred in nearly all rhinocerotids (Prothero et al., 1986; Heissig, 1989; Cerdeño, 1995; Antoine, 2002; Prothero, 2005). Its unique appearance in the members of Aceratheriini and Teleoceratini lies in its position relative to the infraorbital foramen (Cha. 24). In the primitive rhinocerotid *Trigonias* and the extant *R. unicornis*, the nasal notch was shallow, and the infraorbital foramen was behind the notch and above the P4 level (Gregory and Cook, 1928; Prothero, 2005). However, in the Paleogene *Aprotodon* and *Molassitherium*, the nasal notch was moderately retracted, at P3, and the infraorbital foramen was posteriorly moved to below its posterior end. By the Late Miocene, this foramen had increased to three openings in *Shansirhinus* and *Teleoceras* but remained below the ventral edge (Fig. 3) (Qiu and Yan, 1982; Qiu and Xie, 1997; Deng, 2005; Prothero, 2005; Lihoreau et al., 2009).

The broadening and shortening tendencies of the skull are exclusively occurred in Aceratheriini and Teleoceratini and have important systematic value (Heissig, 1989; Cerdeño, 1995). However, both are gradually changing characteristics, and with a long history of evolution; its maximum degree has occurred in only very advanced taxa. Based on the maximum length/maximum width ratio (Cha. 78), several genera have the brachycephalic skull, including *Peraceras, Teleoceras, Brachypotherium*, and *Prosantorhinus* (Prothero, 2005; Heissig, 2017). The brachycephalic skull, accompanied by graviportal limbs, occurs in *Peraceras, Teleoceras*, and *Prosantorhinus* (Prothero, 2005; Heissig, 2017). Both characters are related to a short and massive body and are notable aspects of Aceratheriini and Teleoceratini’s appearance. The graviportal limbs are exemplified in several aspects, and we have added new features to discern, such as the length ratio between the dorsal and lateral sides of the magnum (Cha. 311), the length/width ratio of the third metacarpal (Cha. 329, 330), and the outline of the unciform (Cha. 316) (Fig. 5).

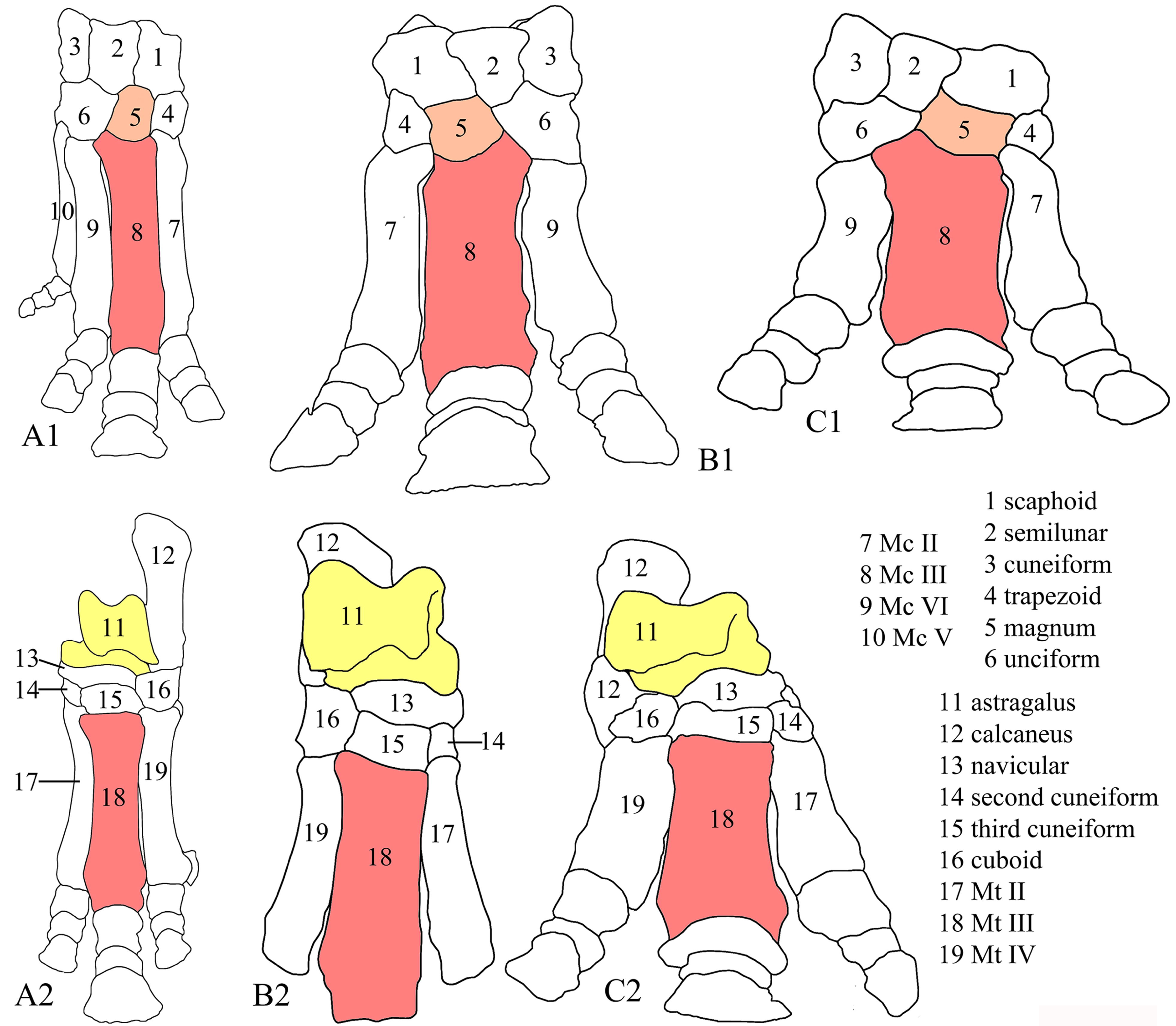

In addition to graviportal limbs, Aceratheriini and Teleoceratini demonstrate another diagnostic character regarding the postcranial bones: the primitive members maintain the functional fifth metacarpus (Heissig, 1989, 1999; Prothero, 2005). The functional fifth metacarpus is inherited from the primitive ancestor of the Rhinocerotidae, and has been reduced to a small sesamoid like bone in the advanced taxa of Aceratheriini and Teleoceratini, such as *Chilotherium* (Fig. 5) (Ringström, 1924; Deng, 2002).

There is only one feature of the cheek teeth (Cha. 209) listed within the synapomorphies of Aceratheriinae, but two other features, which are traditionally considered as a diagnosis of Aceratheriini or Teleoceratini, are not present in the list because they are not apomorphies: deep constriction of both lingual cusps of the upper cheek teeth, with the maximum degree that is just slightly weaker than that of elasmotheres; and the continuous lingual cingulum of the upper premolars, which is inherited from a primitive ancestor (Heissig, 1989, 1999) (Fig. 6). The former could complicate the occlusal pattern of the cheek teeth, and the latter would become the lingual enamel wall of significant worn teeth. Both features are extremely useful for determining the taxonomic identification of the material (Fig. 6).

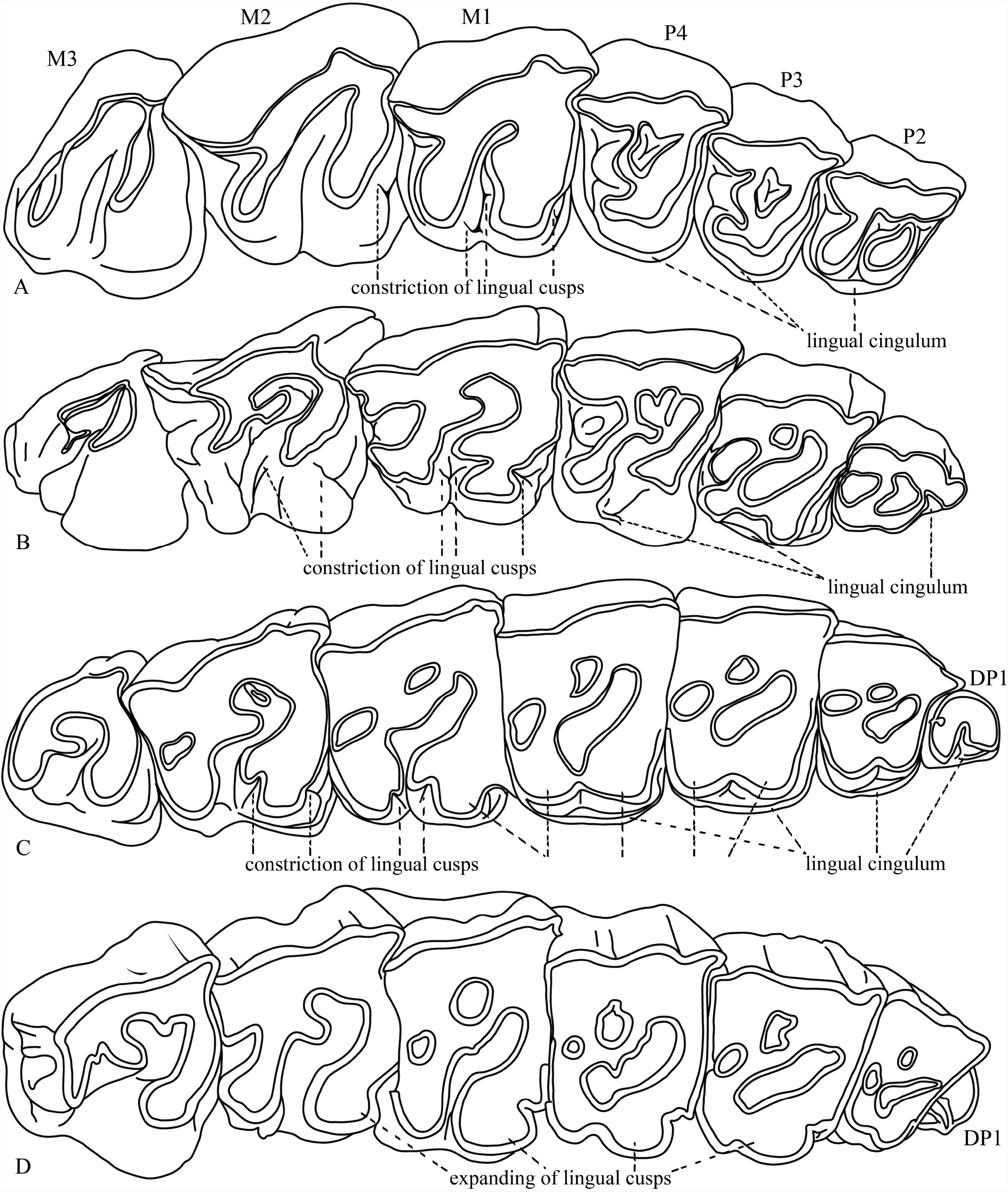

### Phylogenetic relationships of Aceratheriinae

The present analysis found that Aceratheriini and Teleoceratini are closely related in respect to other groups in both the consensus tree and the 50% majority tree, which is consistent with the results of Heissig (1989, 1999) and Cerdeño (1995) (Figs. 1, 2). Here, we use the node-based definition of Aceratheriinae as the last common ancestor of *Aprotodon, Diaceratherium*, and *Aceratherium*, and all of its descendants.

Among the 24 taxa previously referred to as Aceratheriinae, *Turkanatherium* is precluded from this group and recovered as an early diverging taxon of Rhinocerotidae. Deraniyagala (1951) described and established *Turkanatherium* based on materials from the Middle Miocene Moruorot, Kenya, while, Arambourg (1959) and Hooijer (1963, 1966) referred this taxon to *Aceratherium* without further discussion. However, Geraads (2010) re-identified it as *Turkanatherium* based on the differences between *Turkanatherium acutirostratum* and *Aceratherium incisivum*, and attributed it to the subtribe Aceratheriina. The position of *T. acutirostratum* was recently reported within Elasmotheriinae by Geraads et al. (2016), who, however, suggested that their results should be interpreted cautiously. Furthermore, two features of this genus differ from Aceratheriinae; the nasal notch is shallow at the level of P2, and the infraorbital foramen is behind the nasal notch. However, its upper premolars have a metacone rib on the ectoloph and have lost the lingual cingulum, which are similar to Rhinocerotinae but not to Aceratheriinae (Fig. 6). *Turkanatherium* maintains a notable primitive structure on the upper premolars; the metaloph has yet been fully formed. Evidently, *Turkanatherium* is not an acerathere, so we followed the result of the analysis and assigned it as a basal rhinocerotid.

#### Teleoceratini

The Teleoceratini tribe was established based on the genus *Teleoceras* from the Late Miocene in North America, with five widely recognized genera, *Diaceratherium, Brachypotherium, Teleoceras, Prosantorhinus*, and *Aprotodon* (Heissig, 1989, 1999; Prothero and Schoch, 1989; Prothero, 2005). However, the present analysis questioned previous taxonomic opinions and reconstructed a different clade that includes *Diaceratherium, Brachypotherium, Teleoceras, Prosantorhinus*, and *Alicornops*, but precludes *Aprotodon*, which, together with *Mesaceratherium*, was recovered as an early diverging taxon of Aceratheriinae (Fig. 2, Node B). *Aprotodon* from the Late Oligocene in Asia has long been recognized as an early member of Teleoceratini: the skull has a slender outline, the parietal crests fuse and form the sagittal crest, and the upper premolar has yet to be fully molarized (Forster-Cooper, 1915; Borissiak, 1954). In Cerdeño’s (1995) analysis, it was recovered as the most basal Rhinocerotinae. After the description of the new materials of *Aprotodon* from China, Qiu and Xie (1997) likewise suggested it should be an early diverging lineage of Rhinocerotidae. *Mesaceratherium* from the Late Oligocene of Eurasia is the earliest genus of Aceratheriini, with the less molarized premolars (Heissig, 1969). Antoine (2010) placed it as a sensu lato acerathere.

This reference is confusing because the latter has lost the upper incisors I1 and its limb bones are slender. In addition to the synapomorphies listed above, *Alicornops* shows a robust distal limb bone, close to that of Teleoceratini. The lateral and medial sides of the diaphysis of the third metacarpus are irregular, so the diaphysis has no the narrowest position. The third metacarpus is length 90–123 mm, similar to *Diaceratherium aginense* but distinct from *Aceratherium* and *Chilotherium*, in the latter two the narrowest position of the diaphysis is at the proximal extremity and the maximum length ranges from 119 to 112 mm and from 120 to 123 mm, respectively (Répelin, 1917; Hünermann, 1989; Cerdeño and Sánchez, 2001; Deng, 2002). The outlines of the astragalus in *Alicornops* are more flattened than *Chilotherium*, which is the most advanced Aceratheriini (Ringström, 1924; Deng, 2002). These differences in the distal limb bone characteristics suggest that *Alicornops* had evolved toward the graviportal limbs of Teleoceratini, but not Aceratheriini.

#### Aceratheriini

The results of the present analysis did not support the previously established monophyly of Aceratheriini (Prothero and Schoch, 1989; Heissig, 1989; Cerdeño, 1995). *Turkanatherium* has been precluded from Aceratheriinae. The rest of the 17 taxa, except *Mesaceratherium*, which is in a clade with *Aprotodon* in both the strict and 50% majority consensus trees, have been decomposed into seven single genus clades and three small clades (Fig. 2) (Node D, E, F).

In clade of *Hoploaceratherium, Aphelops*, and *Acerorhinus*, the relationship between the former two has been noted in previous studies because of their longer nasal bones (Heissig, 1989). Additionally, among the eight synapomorphies supported node D, the most significant is a uniquely derived feature, the expansion of the lingual cusps of the upper cheek teeth. The constriction of the lingual cusps is an efficient way to complicate the occlusal pattern of the cheek teeth and is present on nearly all lineages of rhinocerotids, excluding the members of Rhinocerotini that expanded the lingual cusps while losing the constriction. The combination of both characters in *Acerorhinus* was first noted by Deng (2000): the expanded and rounded lingual cusps in the upper cheek teeth and the shallow constriction in the upper molars; however, the antecrochet remains present, although with a narrow outline (Fig. 6). This synapomorphy, found only in these three genera, powerfully demonstrates a closer evolutionary relationship (Node D). Another small clade clustering four Late Miocene genera, including *Aceratherium, Peraceras, Chilotherium*, and *Shansirhinus*, represented the most advanced group of Aceratheriini, with a Bremer support value 2 (Node E, Char. 26^0-1^, 42^0-1^, 56^0-1^, 60^0-1^, 183^0-1^, 346^1-0^). Furthermore, this clade has a feature inherited from their primitive ancestor: a very deeply constricted lingual cusp that creates a triangular cusp and large antecrochet, represented an adaptation of evolution of the cheek teeth within Aceratheriinae, distinct from the clade including *Acerorhinus* (Fig. 6).

One unexpected clade is *Galushaceras* and *Subchilotherium*, both of which are from the Middle Miocene (Heissig, 1972; Prothero, 2005) (Node F) (132^1-0^, 154^2-3^, 164^1-0^, 184^0-1^). The latter has long been considered a close relative of *Chilotherium* (Heissig, 1989; Deng and Gao, 2006). In the present analysis, both genera are united mainly based on their less advanced cheek teeth, which have a weak constriction and no crista and medifossette. Additionally, *Galushaceras* has a more advanced nasal notch (P4/M1) than that of *Subchilotherium* (Prothero, 2005). The clade of *Galushaceras* and *Subchilotherium* is closer to the clade including *Aceratherium* regarding the specialized pattern of the cheek teeth: the lingual cusps are constricted and have no expansion. We here combined both small clades together and considered the combination as a sister group of the clade that includes *Acerorhinus* (Fig. 2).

When discussing the phylogeny of Aceratheriinae, Heissig (1973, 1989), Prothero et al. (1986) and Cerdeño (1995) did not distinguish its difference from the tribe Aceratheriini. In the present analysis, the less advanced genera of Aceratheriini were recovered as a set of early diverging clades, each containing a single genus and, included *Molassitherium, Protaceratherium, Plesiaceratherium, Floridaceras, Dromoceratherium*, and *Chilotheridium* (Fig. 2). The first three from the Late Oligocene or the Early Miocene are the early members with assemblages of the primitive features; the skull is slender, the nasal notch is shallow (at the level of P3), the upper incisor I1 and the lower incisor i2 are moderately enlarged, the cingula of the cheek teeth are intact, the lingual cusps of the cheek teeth are moderately constricted, the metapodials are slender, and the carpi and tarsi are narrow and high (Roman, 1912; Yan and Heissig, 1986; Antoine, 2002; Lihoreau et al., 2009; Lu et al., 2016). *Chilotheridium* from Africa in the Early Miocene is also an early diverging genus, which is consistent with its mosaic morphology; it has slender skull and limb bones, but complicated cheek teeth and a horned nasal bone at the subterminal position (Deraniyagala, 1951). *Floridaceras* and *Dromoceratherium* are two moderately specialized genera; their unstable relationships are partly the result of limited available materials (Crusafont et al., 1955; Wood, 1964; Prothero, 2005). Both were recovered as two independent clades in the present analysis.

Evidently, early aceratheres underwent independent adaptations several times, such as the lower incisors i2 of *Aprotodon* and the shorter nasal bone of *Molassitherium*. If we follow the traditional taxonomic framework to assign such taxa in Aceratheriini or Teleoceratini, both tribes will inevitably become a paraphyletic group. Given the results of the cladistic analysis, we reconstructed *Molassitherium, Protaceratherium, Plesiaceratherium*, and *Chilotheridium*, and the clade including *Aprotodon* and *Mesaceratherium* as early diverging basal taxa. Aceratheriini and Teleoceratini are two advanced tribes; the former tends to exhibit a reduced I1 and a further retracted nasal notch, and the latter has an enlarged I1 but graviportal limbs (Fig. 2).

## CONCLUSION

1. According to the results of the cladistic analysis of the newly established matrix, Aceratheriinae is defined as the last common ancestor of *Aceratherium, Diaceratherium*, and *Aprotodon* and all of its descendants (node-based). *Turkanatherium* is not an acerathere.
2. The diagnosis of Aceratheriinae is revised as follows. It is of medium-large size; tended to evolve a brachycephalic skull, and the presence of a nasal horn varies. It has a deep nasal notch, at the level of P3–M1, with an infraorbital foramen below the nasal notch. It exhibits an enlarged or lost upper incisor I1, the lower incisor i2 is tusk-like, even extremely enlarged, and the labial cingulum of the upper and lower cheek teeth are reduced with residuals, but the lingual cingulum of the upper premolars is unreduced; lingual cusps of the upper cheek teeth could be constricted, or sometimes also expanded. It has a functional fifth metacarpus and its limbs have a tendency toward graviportality, the distal parts of which are always massive and flattened.
3. Aceratheriini and Teleoceratini are considered to be two advanced tribes. Teleoceratini includes *Alicornops, Diaceratherium, Prosantorhinus, Brachypotherium* and *Teleoceras*. Aceratheriini includes *Hoploaceratherium, Acerorhinus, Aphelops, Aceratherium, Peraceras, Chilotherium, Shansirhinus, Galushaceras* and *Subchilotherium*. Members of Aceratheriini are divided into two subclades based on the skull outline and the occlusal pattern of the cheek teeth.
4. The clade uniting *Aprotodon* and *Mesaceratherium* is the earliest diverging group within Aceratheriinae, with an extremely enlarged i2. *Molassitherium, Protaceratherium, Plesiaceratherium*, and *Chilotheridium* are also reclassified as a set of basal aceratheres. The phylogenetic positions of *Floridaceras* and *Dromoceratherium* are unstable because of the limited materials, and thus are tentatively reconstructed as basal aceratheres.

## ACKNOWLEDGMENTS

We thank Chen, S. Q. from the Hezheng Paleozoological Museum, Zheng, X. T. from the Tianyu Museum, Wang, S.-Q. from the Institute of Vertebrate Paleontology and Paleoanthropology, Chinese Academy of Sciences. Special thanks go to editors and reviewers for their constructive suggestions for improvement of the manuscript. Mr. Chen, Y., Mr. Xu, Y., and Dr. Sun, D. H. prepared the drawing. This research was supported by State Key Laboratory of Palaeobiology and Stratigraphy (Nanjing Institute of Geology and Palaeontology, CAS) (No. 203113), Key Laboratory of Vertebrate Evolution and Human Origins of Chinese Academy of Sciences (Institute of Vertebrate Paleontology and Paleoanthropology, CAS) (LVEHO19003), Strategic Priority Cultivating Research Program, Chinese Academy of Sciences (XDB26000000, XDA20070203, QYZDY-SSW-DQC022, GJHZ1885).

